# Optogenetically-Induced Population Discharge Threshold as a Sensitive Measure of Network Excitability

**DOI:** 10.1101/641126

**Authors:** D.C. Klorig, G.E. Alberto, T. Smith, D.W. Godwin

## Abstract

Network excitability is governed by synaptic efficacy, intrinsic excitability, and the circuitry in which these factors are expressed. The complex interplay between these factors determines how circuits function and, at the extreme, their susceptibility to seizure. We have developed a sensitive, quantitative estimate of network excitability in freely behaving mice using a novel optogenetic intensity-response procedure. Synchronous activation of deep sublayer CA1 pyramidal cells produces abnormal network-wide epileptiform population discharges (PD) that are nearly indistinguishable from spontaneously-occurring interictal spikes. By systematically varying light intensity, and therefore the magnitude of the optogenetically-mediated current, we generated intensity-response curves using the probability of PD as the dependent variable. Manipulations known to increase excitability, such as sub-convulsive doses (20 mg/kg) of the chemoconvulsant pentylenetetrazol (PTZ), produced a leftward shift in the curve compared to baseline. The anti-epileptic drug levetiracetam (40 mk/kg), in combination with PTZ, produced a rightward shift. Optogenetically-induced population discharge threshold (oPDT) baselines were stable over time, suggesting the metric is appropriate for within-subject experimental designs with multiple pharmacological manipulations.

**Significance Statement:** Abnormal excitability is associated with a number of neurological disorders, including epilepsy. Excitability can be measured in single cells *in vitro*, but it is difficult to extrapolate from these values to the functional impact on the associated network. Epileptiform population discharges are network-wide events that represent a distinct transition from normal to abnormal functional modes. We developed a new method that uses light intensity-response curves to precisely determine the threshold for this transition as a surrogate measure of network excitability.

## Introduction

Seizure thresholds are commonly used to determine excitability and seizure susceptibility in animal models of epilepsy and to assess the effectiveness of therapeutic intervention (Spiegel, 1937; Ziskind et al., 1946; Swinyard et al., 1952; Barton et al., 2001). However, seizures have lasting effects on the brain, including widespread changes in gene expression (Altar et al., 2004), reduced seizure thresholds, and increased seizure severity (Racine, 1972a). This complicates the interpretation of within-subject experiments and limits the ability to make multiple measurements over time. Furthermore, excitability is dynamically modulated by ongoing brain activity and behavioral state, which introduces significant variability, limiting the precision of acute threshold measurements. Single pulse electrical stimulation, a technique used in humans undergoing intraoperative monitoring, provides a measure of network excitability without inducing seizure (Matsumoto et al., 2017). However, this method is limited by the uncertainty of the cells and pathways stimulated (Histed et al., 2009), and the ambiguity of amplitude based LFP measurements in relation to underlying activity (Hales and Pockett, 2014; Herreras, 2016). Excitability and seizure susceptibility can also be estimated by quantification of spontaneous seizures in models of epileptogenesis, but this approach is complicated by the unpredictable occurrence of seizures, requiring constant video-EEG monitoring, prolonged drug administration, and large numbers of animals (Löscher, 2011).

In order to obtain more precise and reliable measurements, we have developed a rigorous approach for quantifying instantaneous network excitability, without inducing seizures, using single pulse light intensity-response curves to determine population discharge thresholds. Optogenetically-induced population discharges (oPDs) are all-or-none, network-wide events with waveforms and latencies similar to spontaneously-occurring interictal spikes (IIS). We show that induced oPDs occur with a higher likelihood with increasing light intensity. By delivering a range of light intensities with randomized presentation order and modeling oPD probability using logistic regression, we account for short-term fluctuations in excitability and derive a new metric, the I50, defined as the intensity of light that produces an oPD with a probability of 0.5. Using established pharmacologic modulators, we demonstrate that the I50 is sensitive to shifts in excitability. The oPDT can be monitored over long timescales with high temporal resolution to fully capture the dynamics of network excitability. Furthermore, because it does not involve the generation of after-discharges (ADs) or seizures, the oPDT can be used as a measure of excitability in a range of normal and pathological conditions other than epilepsy.

Using the same preparation, we show that repetitive stimulation with light of a sufficient intensity produces a robust AD and kindling with daily ADs results in overt behavioral seizure. In the same animal, the optogenetic AD threshold (oADT) and the oPDT are equivalent in terms of light intensity. The oADT, however, requires an additional accumulating process involving multiple oPDs in a short period of time. Comparing the two metrics allows for separation of the contribution of instantaneous excitability, and robustness to seizure, or susceptibility, features that are inseparably intertwined in other measures of seizure threshold.

## Materials and Methods

### Subjects

Male Thy1-ChR2-YFP (founder line 18) transgenic mice (Arenkiel et al., 2007; Wang et al., 2007) (Stock # 007612, The Jackson Laboratory) were used for chronic optogenetic stimulation and recording (n = 35). Thy1 line 18 mice express wildtype ChR2 fused to EYFP in area CA1, subiculum, and layer 5 of cortex (Arenkiel et al., 2007). In area CA1, expression is specific to excitatory pyramidal neurons and is concentrated in calbindin-negative cells in the deep pyramidal cell sublayer (Dobbins et al., 2018). These mice can be bred as homozygotes, ensuring consistent expression levels and patterns from animal to animal, a significant advantage for reproducibility. For this reason, they are used in a number of optogenetics studies (Ting and Feng, 2013). Not all animals were used for all experiments; exact numbers are reported throughout. All animal experiments were approved by the Institutional Animal Care and Use Committee (IACUC) of Wake Forest University in agreement with National Institutes of Health and United States Department of Agriculture guidelines.

### Chronic Implant

Details of the implantation surgery and chronic recording array are described in a previous publication (Klorig and Godwin, 2014). Briefly, Thy1 mice (age 2-9 months, mean 120 ± 9 days) were anesthetized with isoflurane and placed in a stereotaxic device. All surgical procedures were performed under red light to avoid sustained activation of ChR2. Metal ground screws were secured to the cranium bilaterally so that they were in contact with the subarachnoid space posterior to the transverse sinus. A total of 8 tungsten microwires were implanted in a satellite array in cortical, subcortical, and hippocampal locations: prefrontal cortex, (PFC, AP: 5.58, L: 0.37, DV: −1.81, in mm, from the interaural plane), anterior medial thalamic nucleus (AMTh, AP: 2.86, L: 0.24, DV: −3.62), dentate gyrus (DG, AP: 1.50, L: 1.39, DV: −1.61), hippocampus (CA3, AP: 1.34, L: 2.88, DV: −2.12), entorhinal cortex (Ent, AP: 0.19, L: 3.90, DV: −2.73, angled 16° posterior), subiculum (sub, AP: 0.88, L: 1.02, DV: −1.43), and left CA1(AP: 1.00, L: −2.36, DV: −1.20). (Fig. 1A). A magnetic rotary fiber connector with an attached microwire (optrode) was placed in intermediate CA1 near subiculum (CA1a) (AP: 1.00, L: 2.36, DV: −1.20, from the interaural plane). The stimulation fiber was a multimodal fiber with a 200 µm diameter core and a numerical aperture of 0.39 (FT200UMT, ThorLabs). The recording electrode extended 0.5 mm beyond the fiber tip. The optical fiber was situated 0.4 mm dorsal to the pyramidal cell layer and 0.3 mm dorsal to the basal dendrites. The optrode was configured to minimize the photovoltaic effect (<0.02% LFP amplitude) by cutting the electrode at an angle so that the exposed metal surface was not directly illuminated by the implanted fiber (Cardin et al., 2010). Wild-type (n = 2) and post-mortem control (n = 4) recordings were used to confirm the absence of artifact. This setup allowed for high quality recordings, free of movement artifacts, even during behavioral seizures.

**Figure 1.**
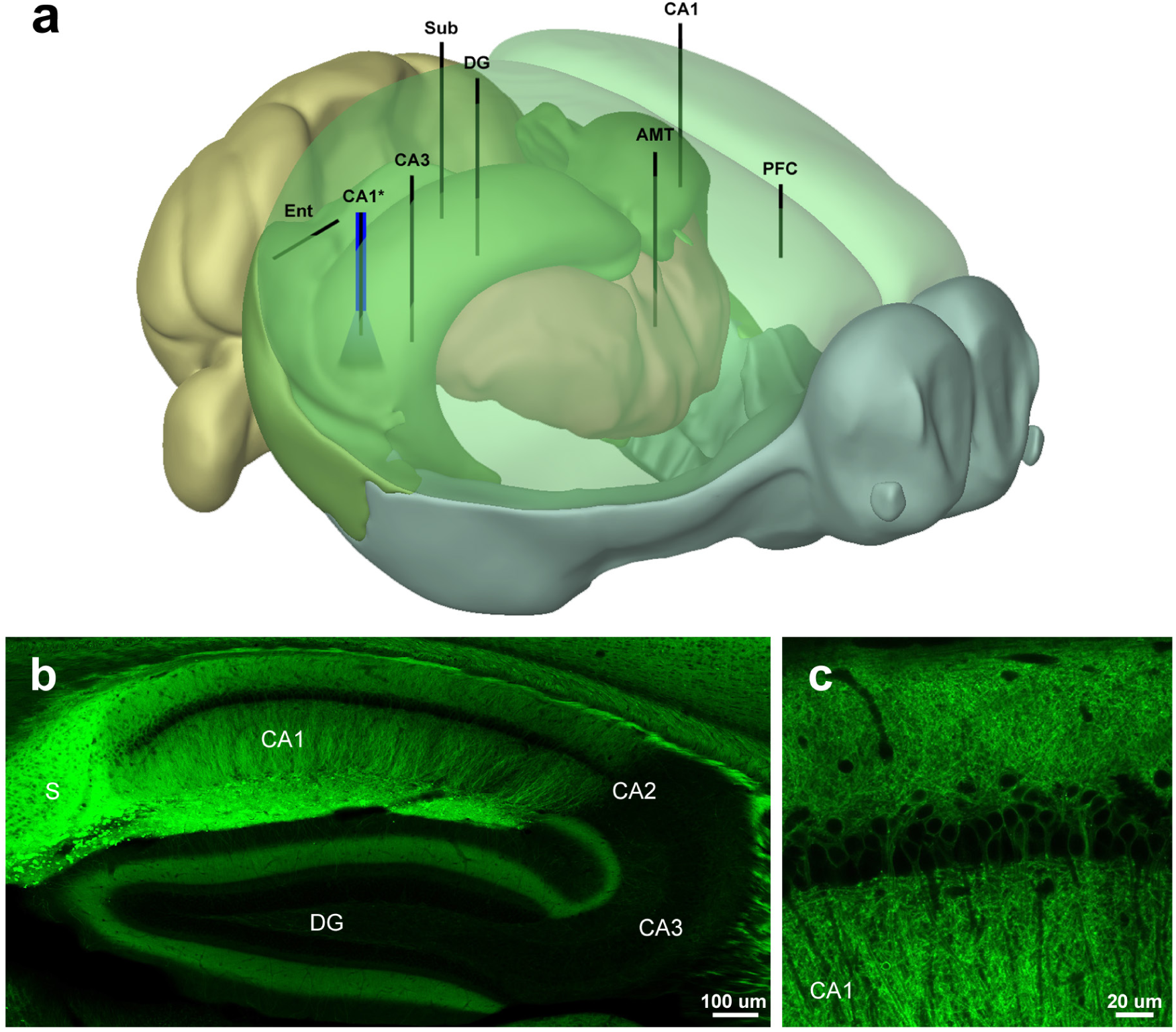
Electrode placement and expression patterns in the Thy1-ChR2 (line 18) mouse. (**a**) Schematic showing the placement of the electrode array and optrode. (**b**) Distribution of ChR2-EYFP in the hippocampus of the Thy1 (line 18) mouse. Note high expression levels in CA1 and subiculum. 10x tiled confocal image. (**c**) ChR2-EYFP is expressed in deep pyramidal neurons in CA1 (Dobbins et al., 2018). 63x oil immersion confocal image. The 3D model was generated using Brain Explorer Software courtesy of the Allen Brain Institute (http://mouse.brain-map.org/static/brainexplorer).

### Light stimulation and recordings

Mice were allowed to recover 1 week prior to recordings. Wideband (0.3-20kHz, sampled at 40kHz) depth recordings were made using the SciWorks recording system (DataWave Technologies), AM-3600 extracellular amplifiers (A-M Systems), and T8G100 headstage amplifiers (TBSI). Photostimulation was performed with a fiber-coupled LED with peak emission at 470 nm (M470F3, ThorLabs). LED intensity was controlled using an LED driver with analog modulation (LEDD1B, ThorLabs). Analog output pulses were generated within SciWorks and used to control the LED driver. Light pulses were continually monitored with a silicon photodiode placed near the emitter. A custom op-amp based current-to-voltage converter circuit was used to linearize the photodiode output and the resulting signals were recorded along with the electrophysiology data. Two measures of light output are provided; the power (mW) at the tip of the source fiber and an arbitrary linearized scale. Light power ranged from 0.43 – 4.15 mW at the fiber tip corresponding to an irradiance of 0.2 – 1.96 mW/mm^2^ at the recording site estimated using the empirically derived model described in (Aravanis et al., 2007; Yizhar et al., 2011).

### Video and Behavioral Scoring

Recording sessions were video recorded and an infrared LED synchronized to the stimulus onset was placed in view of the video camera for precise temporal alignment between the video and the recordings. Video was used for assessment of behavioral seizures using a modified Racine scale (Pinel and Rovner, 1978; Racine, 1972b). Latencies to seizure stage relative to the onset of the stimulus and durations were also recorded.

### Histological Verification of Electrode Placement

On completion of recording experiments, mice were anesthetized with Euthasol (Virbac Animal Health), electrolytic lesions were performed at each electrode site (20 nA, 10 sec), then animals were transcardially perfused with saline and 4% paraformaldehyde solution in PB. After removal and post-fixing, brains were sectioned in 50 µm slices using a vibratome. The slices were then mounted on slides and stained using cresyl violet (Nissl). Each slice was imaged using a light microscope and the stereotaxic coordinates of the implanted electrodes were recorded. Animals were excluded from further analysis if electrodes utilized in those analyses were located outside of the target areas (n = 5).

### Confocal Microscopy

ChR2-eYFP expression in Thy1 mice was imaged in non-implanted animals. Confocal imaging was performed on a Zeiss LSM 710 confocal microscope with 10X, 20X air and 60X water immersion objectives.

### Experimental Design and Statistical Analysis

Data analysis was performed using custom scripts written in MATLAB (MathWorks). The suite of stimulus presentation and analysis tools used to generate and calculate the oPDT is available on github (https://github.com/neuroptics/optoDR).

### Optogenetic Population Discharge (oPD) Intensity-Response Curve

Single square pulses (10 ms) of light were delivered at a range of intensities (20 levels, 0.43 - 4.15 mW at fiber tip, 0.2 – 1.96 mW/mm^2^ at the recording site). Sixty repetitions of each intensity were presented in randomly ordered blocks with a 1 s or 3 s interval between pulses. A timing pulse was used to precisely align responses to the onset of the LED. For the population discharge curves, time-locked evoked responses were extracted, the 2^nd^ derivative calculated, and a sliding window (10 ms) was used to calculate root-mean-square (RMS) power via convolution. The max RMS within a specified time window was calculated, sorted by magnitude, and plotted for each channel over all stimulation levels. The channel with the sharpest transition between sub-PD and PD was chosen and the threshold set (typically contralateral CA1, but occasionally DG or CA3 were chosen). PD detection was verified visually. All subsequent analyses for a given animal were performed with the same channel and threshold. The ratio of PD events per level was calculated to generate a probability curve. This curve was fit by the Hodgkin and Huxley formulation of the Boltzmann distribution:

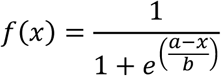

where a is the I50, and b is the slope (Hodgkin and Huxley, 1952), using nonlinear regression (least squares) in MATLAB. The I50 value for each session was then used for further comparisons.

### Optogenetic After-Discharge Threshold (oADT) and Kindling Procedure

oADTs were determined by presenting optical stimulation trains at 6.67 Hz (10 s duration, 4 ms pulse, 150 ms interval, 66 pulses), then increasing the intensity (with a 2 min interval) until an AD was evoked (characterized by high amplitude, self-sustaining activity that persisted > 5 s after the stimulus ended). A modified optogenetic kindling procedure was used that allowed for estimation of oAD thresholds, involving repeated daily light stimulation at increasing intensity until an AD was evoked (with intensities identical to that performed on the first day), similar to established procedures (Bragin et al., 2002; Pinel et al., 1976). ADs were defined as sustained high-amplitude spike and poly-spike activity lasting at least 5s following termination of the stimulus. oAD durations were measured from the start of the stimulus to the last spike of the oAD. Only one oAD was evoked per 24 hour period. This procedure was repeated each day for 15 days.

### Pharmacology

All experiments utilized a within-subjects design. Treatments were compared to a pre-treatment baseline using repeated measures one-way ANOVA with Tukey’s test for multiple comparisons. Saline controls were also performed and compared to pre-injection baselines.

### Statistics and Measures of Reliability

Statistical testing was performed with MATLAB and Prism (GraphPad). All means are reported ± SD, with 95% confidence intervals (CI) where appropriate. Data sets were tested for normality using the omnibus K^2^ test (D’Agostino et al., 1990). Comparisons were performed using repeated measures one-way ANOVA with Tukey’s test for multiple comparisons, paired t-tests, Fischer’s exact test, or Wilcoxon matched pairs test where appropriate. Correlation coefficients were calculated using Pearson’s or Spearman’s method where appropriate. All reported statistics are labeled with the test used. Box and whisker plots have min-max whiskers, 25^th^ to 75^th^ percentile boxes, and the central line is the median. The probability of oAD given at least x oPDs was calculated using the formula: 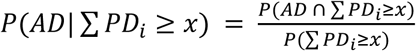. For assessment of oAD probability given oPD on a given stimulus pulse n, we used the formula for conditional probability: 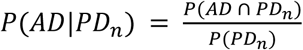. For assessment of oPD_I50_ probabilities given the results of previous trials we used the formula: 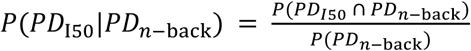. Fisher’s exact test was used to compare proportions.

## Results

In order to explore network propagation of optogenetically induced seizures and PD activity, we developed a multi-site satellite array system consisting of individually placed microwires in perihippocampal structures and an optrode above CA1. Targets included prefrontal cortex (PFC), anterio-medial thalamus (AMTh), entorhinal cortex (Ent), dentate gyrus (DG), subiculum (Sub), hippocampal area CA3 (CA3), and area CA1 bilaterally (CA1-L and CA1-R) (**Fig. 1a)**). These recording sites provide coverage of areas demonstrated to be strongly activated by optogenetic stimulation of CA1 using fMRI (Weitz et al., 2015). Our preparation yielded high quality chronic recordings of multiple interconnected areas, allowing us to assess the impact of optogenetic stimulation on the network. By varying the intensity of light stimulation, we measured AD thresholds using train stimuli (oADT), and the unitary population discharge threshold using single pulses (oPDT). We present the results of the oADT experiments first in order to provide context for the oPDT, but it should be noted that prior ADs, and/or behavioral seizures exposure, are not required to obtain oPDT measurements.

### Measuring the oADT and opto-Kindling

Optogenetic ADs and behavioral seizures were induced using rhythmic squarewave optical stimuli (6.67 Hz, 10 ms pulse, 10 sec train). 6.67 Hz was the lowest frequency that reliably produced an oAD and the longer period (150ms) provided sufficient time between pulses to observe the propagation of induced activity throughout the network (**Fig. 2b**). Stimulus trains were presented every 2 minutes at increasing light intensity until an oAD occurred (**Fig. 2a-c**). Subthreshold stimulation produced local activation at the stimulation site (CA1), but the evoked activity did not spread to downstream areas (**Fig. 2a**, top inset). The mean oAD threshold was 2.11 mW (SD 1.12, 95% CI [1.08-3.15], *n*=7).

**Figure 2.**
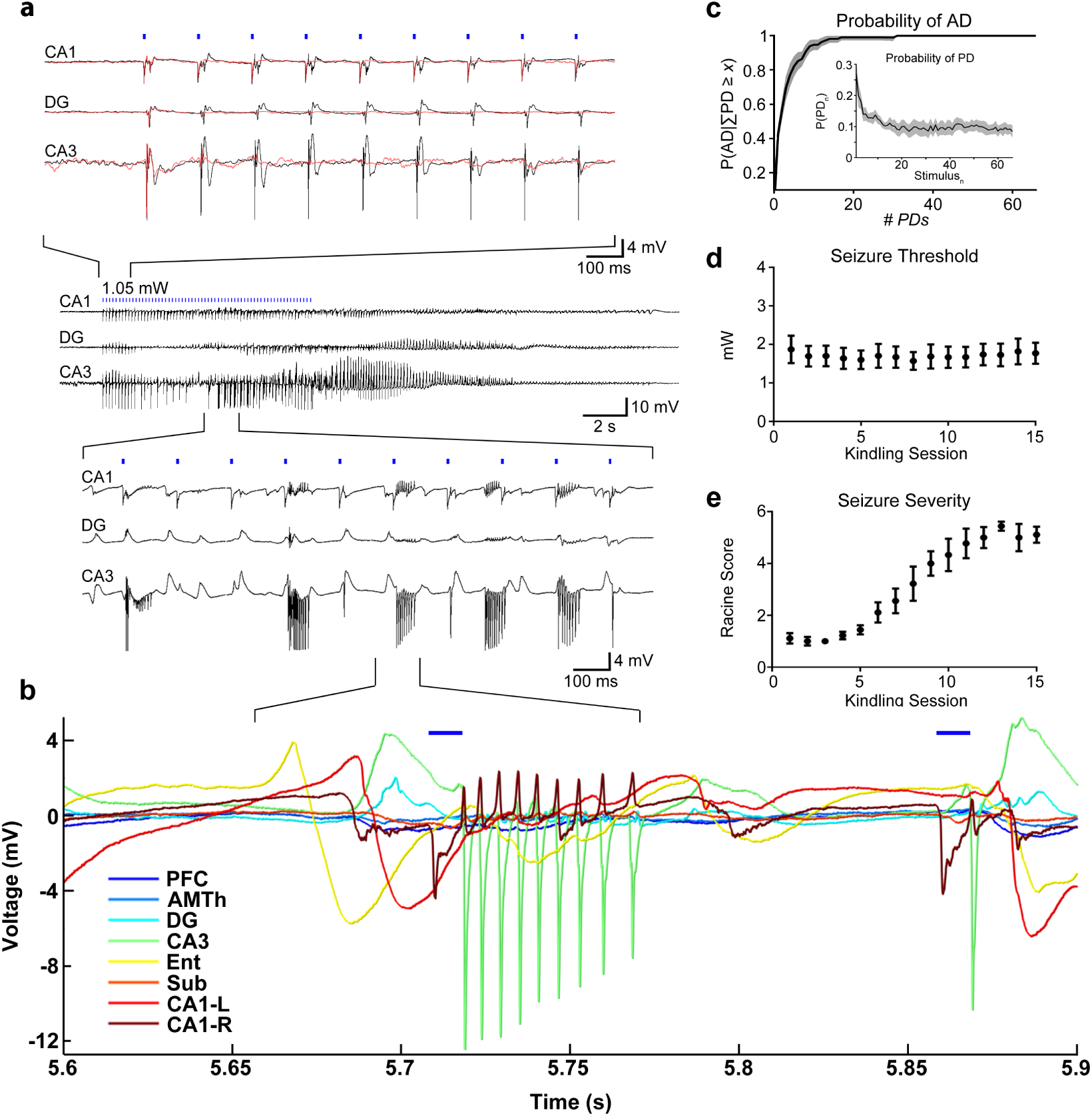
Optogenetic train stimulation produces an AD. Kindling with daily oADs results in an overt behavioral seizure. (**a**) Representative example of an optically-evoked AD. Top Traces: Suprathreshold stimulation levels (for AD) produce network wide PDs (black traces). At subthreshold intensity levels, activation was observed only at the stimulation site (CA1-R), (red traces). Middle Traces: Zoomed out view of oAD. Bottom Traces: Pathological high frequency oscillations (pHFOs) during oAD induction. (**b**) oADs involve pHFOs across multiple sites with a non-zero phase lag. (**c**) The probability of AD given at least x PDs occurring during the stimulus train. Inset: The overall probability of a PD on each pulse of the train, including subthreshold stimuli. Probabilities were calculated for each animal and the mean is displayed with 95% CIs, n = 7 animals. (**d**) Seizure threshold remained stable throughout the kindling period. (**e**) Behavioral seizure severity increased with each AD and was correlated with stimulation number.

Our preparation is configured to stimulate CA1 with minimal damage by situating the optical fiber just above CA1 in visual cortex. In addition to strong expression in CA1, Thy1 mice also express ChR2 in layer 5 of cortex, raising the possibility that these cells were activated by light backscatter. To verify that CA1 is the relevant site of stimulation for seizure induction, we performed control experiments (n = 2) in which the fiber was placed superficially in cortex (0.5mm dorsal to the fiber location in the standard array). Much higher light intensities were required to induce ADs in these animals (16.58 mW and 27.86 mW using a fiber coupled laser), levels sufficient to produce suprathreshold activation of CA1 from this distance, suggesting that activation of cortex is not sufficient to generate ADs and that CA1 is the primary site of activation.

While suprathreshold stimulation produced network wide oPDs following each pulse, high-frequency high-amplitude bursts, and oAD, occasionally the stimulation intensity just below threshold produced synchronous discharges similar to oPDs on the first few pulses (**Fig. 2a**, red traces, top). Likely a result of ChR2 desensitization (Lin, 2010), the probability of an oPD decreases with pulse number (**Fig. 2c** inset), allowing us to estimate the number of oPDs required for an oAD. We found that the probability of oAD depends on the number of oPDs generated by the stimulus (**Fig. 2c**), P(AD) > 0.90 for *x* = 8 oPDs and P(AD)>0.99 for *x* = 17 oPDs (n = 841 stimulus trains). This suggests that the oAD is a result of the accumulated effect of closely spaced oPDs, through some unknown mechanism possibly related to the accumulation of extracellular potassium (de Curtis et al., 2018). oADs occurred during and immediately after stimulation (**Fig. 2a-c**), rather than after a delay, as previously observed with optogenetic seizure induction in motor cortex (Khoshkhoo et al., 2017). The mean oAD duration was 23.1 s (SD 4.84, 95% CIs [22.2, 24.0], *n* = 105 oADs).

Pathological high frequency oscillations (pHFOs, 80-500 Hz) have been detected preceding seizures in a number of animal models and in intracranial recordings from humans with epilepsy (Bragin et al., 1999; Staba et al., 2002; Zijlmans et al., 2012). In the majority (88%, *n* = 105 oADs) of optogenetically induced oADs, pHFOs occurred in CA3, DG, and CA1 (**Fig. 2a and b**). A non-zero phase shift was observed between these areas, suggesting synaptic or ephaptic transmission, rather than volume conduction (Shivacharan et al., 2019) (**Fig. 2b**). The mean latency from the start of the stimulation to burst onset was 6.42 s (SD 2.75, 95% CI [5.87-6.97], *n* = 97 oADs). Epileptiform bursts ranged in frequency from 150 to 400 Hz (6.6 to 2.5-ms intraburst intervals) and were detected in all animals tested (7/7). These bursts were similar (in terms of amplitude and frequency) to patterns previously observed in perforant path-stimulated or kainate-treated rats (Bragin et al., 1997, 2004, 2005).

### Opto-Kindling and the Emergence of Behavioral Seizures

Using this thresholding procedure, we took a subset of animals (n=7) through an opto-kindling procedure based on those performed previously with electrical stimulation (Goddard et al., 1969; Racine, 1972a) (**Fig. 2d,e**). Threshold oADs were presented daily for 15 days. Traditional electrical kindling results in reduced thresholds and increased behavioral severity (Racine, 1972a). In contrast, the oAD threshold remained stable (uncorrelated with presentation #) over the course of kindling (r = 0.0865, p = 0.759, ns, *n*=7) (**Fig. 2d**), in agreement with previously reported optogenetic seizure thresholds obtained by varying stimulus number (Khoshkhoo et al., 2017). The mean oAD threshold during kindling was 1.99 mW (SD 0.827, 95% CI [1.22-2.75], *n*=7). Repeated oAD induction was correlated with increasingly severe behavioral seizures as quantified using a modified Racine scale (r = 0.963, p < 0.0001, *n*= 7) (**Fig. 2e**). Animals progressed to stage 5 seizures after 9.6 days (SD 2.4, 95% CI [7.4-12], *n* = 7), considerably faster than corneal or hippocampal kindled mice (Rowley and White, 2010; Stover et al., 2017), but slower than kindling of piriform cortex (McIntyre et al., 1993). The stereotypical behavioral expression (semiology) of the seizure was similar to that observed with electrical kindling (Kairiss et al., 1984; Racine et al., 1972). Spontaneous seizures were not observed during or following the 15 day kindling period.

### oPD Induction and Threshold Measurement

Given the close relationship between the appearance of multiple network wide oPDs following stimulation and subsequent oADs, we sought to determine if the oPD itself was a sufficient indicator of the oADT and therefore, a potential measure of oAD threshold without the need for oAD induction.

Single pulse optogenetic stimulation of CA1 at a sufficient intensity (mean 2.85 mW, SD 0.248, 95% CI [2.34-3.36], *n* = 30 mice) produced nearly synchronous PDs across the hippocampal and perihippocampal network (**Fig. 3a,b**). The largest amplitude responses were observed in DG and CA3, as well as contralateral CA1. These unitary oPDs closely resembled those induced in the first few pulses of the optical stimulus train used in the oADT procedure, as well as spontaneous type-2 IIS, observed in AD exposed animals (**Fig. 3a**).

**Figure 3.**
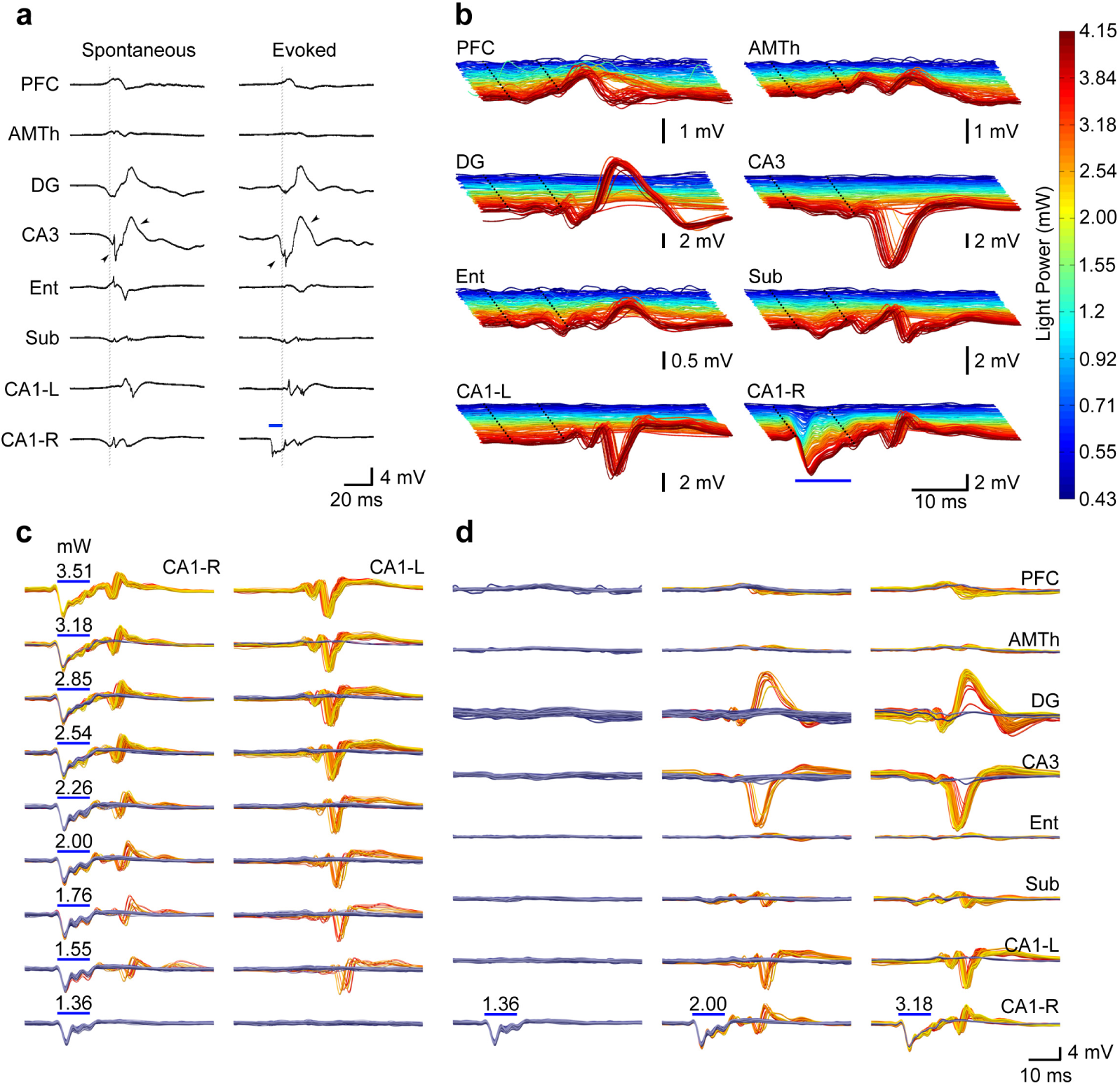
Optogenetic stimulation produces network wide PDs that closely resemble spontaneous IIS with a probability that increases with light intensity. (**a**) Example comparing spontaneous IIS to oPDs. Spontaneous IIS are characterized by near synchronous activation in DG, CA3, and CA1-R, and delayed activation in CA1-L. Evoked discharges are led by activation in CA1 (stimulation site) followed by near synchronous activation of DG, CA3, and CA1 with a latency of ∼10 ms. Note similar timing and wave shape between stimulus evoked and spontaneous discharges (arrow heads). Blue bars indicate onset and duration of light stimulus in CA1. Inter-stimulus interval (ISI) = 3s (**b**) Evoked network activity produced by optogenetic stimulation of area CA1 (right side) at varying light intensities in a representative animal (different from that depicted in **a**). Note large amplitude oPDs evoked at higher intensities. Dotted lines indicate the beginning and end of the light pulse. **(c-d)** Probability of oPD occurrence in downstream sites, rather than amplitude, increases with light intensity at the stimulation site. In area CA1-L, a longer latency response is accompanied by a high amplitude PD. The gold colored traces are those in which a PD was detected, the blue are traces where it was absent. Each plot is an overlay of 60 trials and the color gradient indicates chronological order (dark blue/red for early trials, light blue/yellow for late trials). oPD occurrences: 0/60 @ 1.36 mW, 11/60 @ 2 mW, 58/60 @ 3.18 mW. ISI = 3s. Scale bars in **d** also apply to **c**.

In order to precisely measure the oPDT, we developed a novel optogenetic intensity-response curve procedure that takes advantage of the all or none transition between the sub and suprathreshold response at recording sites downstream of the optogenetic stimulation to accurately detect the occurrence of oPDs (**Fig. 3**). Repeated presentation of the intensity-response curve revealed considerable variability in the response (oPD vs. no oPD) given a particular stimulus intensity (**Fig. 3 c, d**). However, we found analyzing the probability of an oPD given stimulus intensity x, P(oPD|I_x_), allowed for a more robust estimation of threshold.

To generate the oPD probability curve we used twenty intensity levels presented in repeated blocks. Stimuli were presented in random order without replacement so that each stimulus intensity level was presented once per block. Stimulus order was independently randomized for each block. oPD probability was then estimated using multiple replications of stimulus blocks. The intensity levels were selected to linearize the optogenetic response as measured by the mean amplitude of the first peak at the stimulation site, likely a reflection of the interaction between the ChR2 mediated current and dendritic voltage-gated Na^+^ currents (**Fig. 4a-d**). Network oPDs were detected by thresholding a sliding window root-mean-square (RMS) of the 2^nd^ derivative of the peri-stimulus activity (**Fig. 4e-g**).

**Figure 4.**
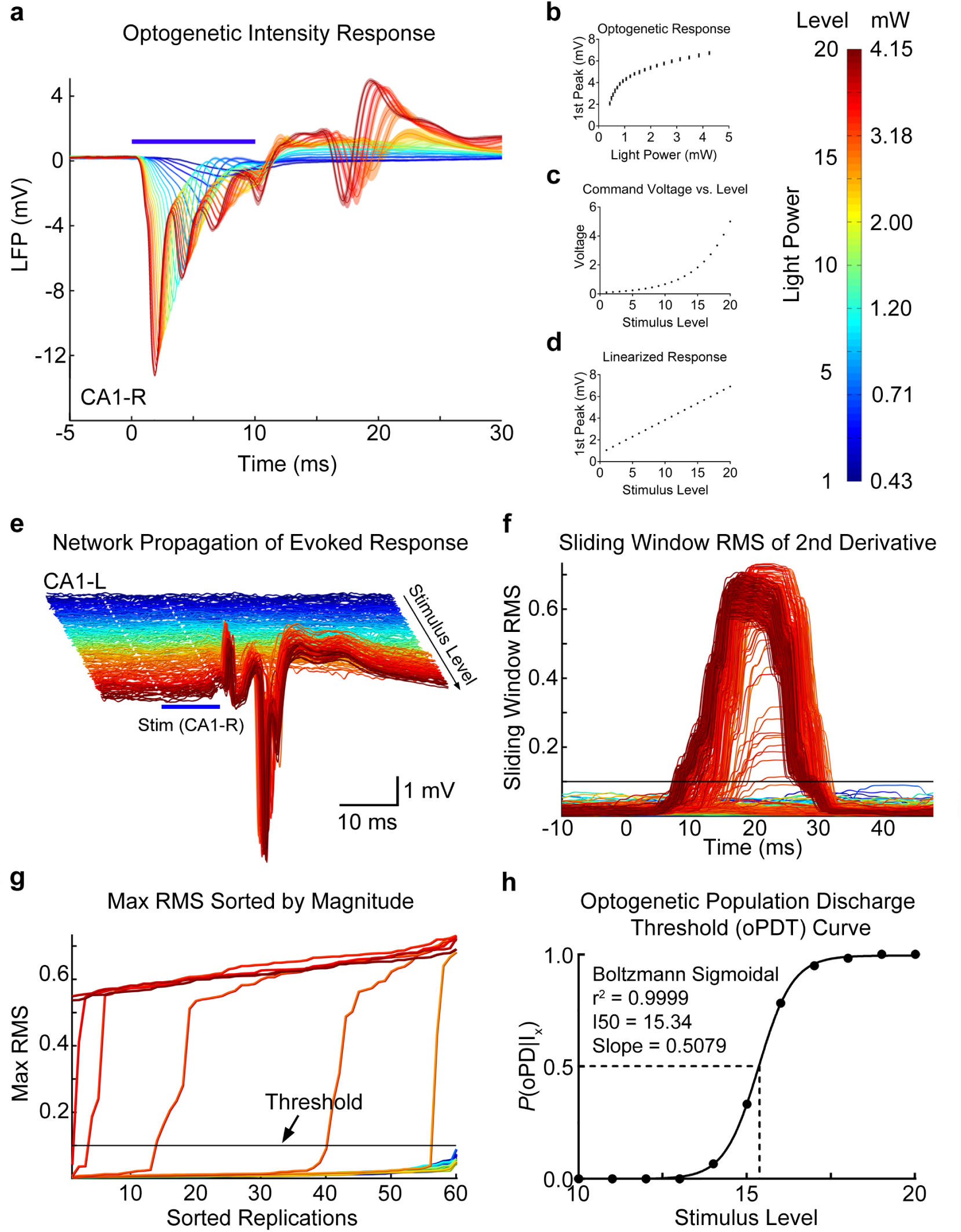
Summary of the oPDT procedure. (**a**) Average LFP response to optogenetic activation in CA1 at varying light intensities. Shaded bars = 95% confidence intervals. (**b**) The short latency LFP response was used as an approximation of ChR2 mediated current to generate a light intensity-response curve. Mean and SD are displayed. (**c**) A series of light intensities was chosen to produce a linearized optogenetic response. (**d**) Linear stimulus response curve. (**e**) The network propagation of evoked activity from the stimulation site (CA1-R, blue bar) to CA1-L. Individual replicates plotted as bandpass filtered (5-300Hz) traces. Colors correspond to the level of the light stimulus (shown in mW on colorbar). 20 levels x 60 replicates = 1200 total stimuli. ISI = 3s. Note high amplitude PDs. (**f**) Sliding window RMS of the 2nd derivative calculated from **e**. Horizontal line indicates the threshold. (**g**) Max RMS (within time window) for each light level sorted by magnitude. Threshold is set in the steep part of the curve. (**h**) oPD ratio (oPDs/total stimuli) plot fit by the Boltzmann sigmoid allows for calculation of an I50, or the LED power at which 50% of the stimuli produce a oPD.

The oPD occurs across the network, and therefore, any channel or combination of channels can be used for oPD detection, including the long latency response at the stimulation site; however, we found that contralateral CA1 provided the greatest separation between sub- and suprathreshold responses. The sub-supra separation was visualized by plotting the sorted peak values of the RMS for each level (**Fig. 4g**). Good separation produces a steep transition with few intermediate values. This plot is also useful for setting the threshold and the steepness of the transition reduces the detection bias introduced by threshold selection. As a result, the oPDT curve is robust to any choice of threshold within the steep transition region of the sorted peak RMS plot.

Once oPDs have been detected, the proportion of oPDs to total stimulus presentations at each level can then be plotted to generate an oPD threshold curve (oPDT). We then fit the data with the Hodgkin and Huxley formulation of the Boltzmann distribution and calculated the I50, or the intensity at which the probability of oPD was 50% (Hodgkin and Huxley, 1952)(**Fig. 4h**). Expressing the I50 in the arbitrary linearized scale (1-20 levels) is convenient for making comparisons, however, it can easily be converted into power (mW) or irradiance (mW/mm^2^) (**Fig. 4b-d**). Although the linearized scale improves the fit of the Boltzmann, substituting mW also produces a sigmoidal curve. Using an arbitrary scale and baseline normalization allows for comparisons between animals even when the relationship between power output and optogenetic response varies. In our experience, the variability in baseline I50 between animals, SD = 3.41 levels (n = 20 animals), was larger than the mean I50 variability within animals, mean SD = 1.10 levels, (n = 3-10 baselines per animal). Since ChR2 expression patterns are preserved between animals, it is likely that between-animal differences are due to variations in the surgical preparation and the transmission efficiency of the implanted fiber.

### Relationship between the oPDT and oADT

Our analysis of the oADT revealed that multiple oPDs must be evoked by the train stimulus in order to generate an oAD (**Fig. 2c**). We compared the oADT and the oPDT measured in the same animals using the same chronically implanted fiber. We found that the oADT was correlated with pre-kindling I50 across animals with a slope approaching 1 (1st Seizure: Lin. Reg. Slope: 1.05, 95% CI [0.677, 1.42], Pearson’s r = 0.956, p = 0.00078, *n* = 7; Avg Seizure: Lin. Reg. Slope: 0.830, 95% CI [0.462, 1.20], Pearson’s r = 0.933, p = 0.0021, n = 7) (**Fig. 5a**). This finding combined with the fact that train stimuli subthreshold for the oPD also failed to produce an oAD, indicate that the oPDT and the oADT are equivalent in terms of light intensity delivered from the same implanted fiber, because both require the oPD.

**Figure 5.**
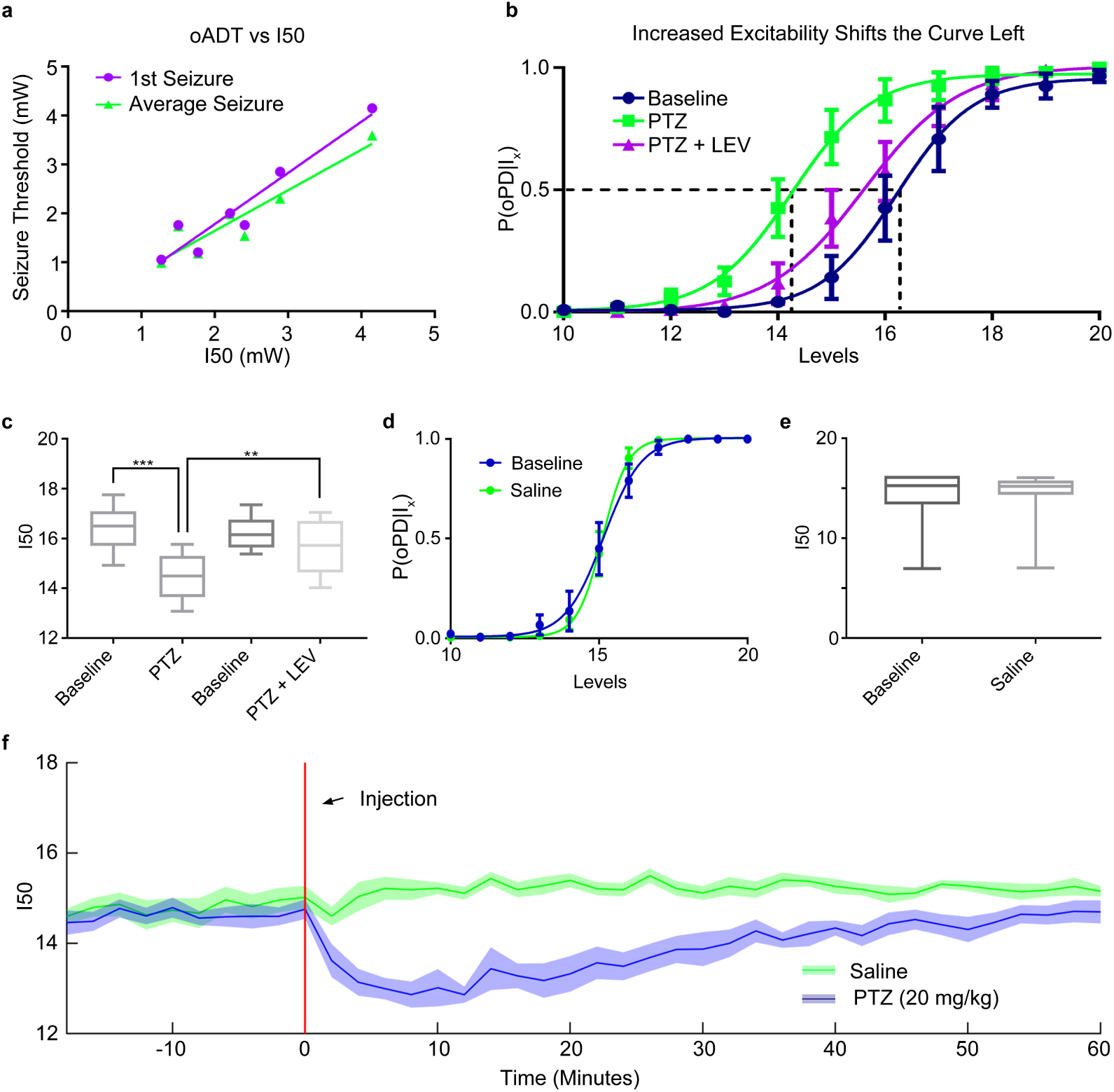
The I50 is sensitive to changes in network excitability. (**a**) The relationship between pre-seizure I50 and the 1st oADT (purple) or the average oADT throughout 15 kindling sessions (green). (**b,c**) The GABAa antagonist PTZ (20 mg/kg) shifts the oPD curve leftward and treatment with the anticonvulsant drug levetiracetam (40 mg/kg) partially reverses the shift. n = 6 animals. *** p = 0.00043, ** p = 0.0034. Curve error bars indicate the 95% CI. (**d,e**) Handling and IP saline administration did not significantly shift the oPD curve compared to baseline (paired t-test, n = 7). (**f**) I50 measurements reveal the time course of PTZ-induced hyperexcitability (disinhibition). Plot of the I50 over time. 6 block bins, n = 9 presentations of each condition (3 trials x 3 animals). Shaded area is mean ± 95% CI.

### Changes in Excitability Shift the oPD Curve

To test the hypothesis that the P(oPD|I_x_) changes with the balance of excitation and inhibition, we used a well-known chemoconvulsant drug, pentylenetetrazol (PTZ), in non-kindled animals. Despite its ability to induce acute seizures, PTZ does not produce a shift in the electrical AD threshold (Karler et al., 1989). In contrast, the oPDT was sensitive to PTZ at sub-convulsive doses (20 mg/kg). The I50 was significantly shifted to the left relative to baseline (Baseline = 16.4 levels, SD 0.943, 95% CI [15.4, 17.4], PTZ = 14.5 levels, SD 0.949, 95% CI [13.5, 15.5], p = 0.00041, *n* = 6) (**Fig. 5a,b**), which indicated a reduction in threshold.

Although the antiepileptic drug levetiracetam (LEV) failed to show efficacy in the traditional MES and s.c. PTZ acute seizure models (Löscher, 2011), it does increase the ADT in kindled rats (Löscher et al., 2000). Pre-treatment with LEV (40 mg/kg) partially reversed the leftward shift produced by subthreshold PTZ and returned threshold to near baseline levels (PTZ+LEV = 15.7 levels, SD 1.11, 95% CI [14.5, 16.8]) (Baseline vs. PTZ, p = 0.00043, Baseline vs PTZ+LEV, p = 0.12, PTZ vs. PTZ+LEV, p = 0.0034, *n* = 6 repeated-measures ANOVA with Tukey’s correction for multiple comparisons) (**Fig.5a,b**). Saline injection did not significantly shift the curve (Baseline vs. Saline, p = 0.99, *n* = 7, paired t-test).

By breaking the stimulus period into bins of presentation blocks, P(oPD|I_x_) can be measured over time. Administration of subthreshold PTZ (i.p.) produced a rapid shift in excitability that partially recovered by about 60 min post-injection (**Fig. 5f**), compared to saline, consistent with the pharmacokinetic curve of PTZ (Yonekawa et al., 1980). For this example, 6 block bins were used to estimate the I50 at 2 min intervals (6 blocks x 20 stimuli per block x 1s ISI = 120s), but higher temporal resolution is possible by decreasing the bin size and/or the number of stimuli per block.

### Stability and Reliability of the oPDT

We assessed the reliability of oPDT measurements, in terms of short- and long-term stability, by analyzing baseline data. Two inter-stimulus intervals (1s and 3s) were used to produce intensity-response curves. This allowed for a comparison of the stability of baseline I50 values generated using each protocol, with the caveat that the data came from different animals. Baselines were selected for further analysis if they were collected in naïve animals (pre-kindling, pre-treatment) (*n* = 67 sessions in 13 animals for 1s, *n* = 41 sessions in 17 animals for 3s int).

First we looked at the test-retest reliability of measurements (20 levels, 60 replications each) taken on consecutive days (**Fig. 6a).** The measured I50s were consistently reliable with an overall correlation coefficient > 0.85 (1s int: r = 0.917, 95% CI [0.705, 0.979], p < 0.0001, *n* = 11 pairs, 3s int: r = 0.856, 95% CI [0.446, 0.969], p = 0.0032, *n* = 9 pairs, Pearson’s).

**Figure 6.**
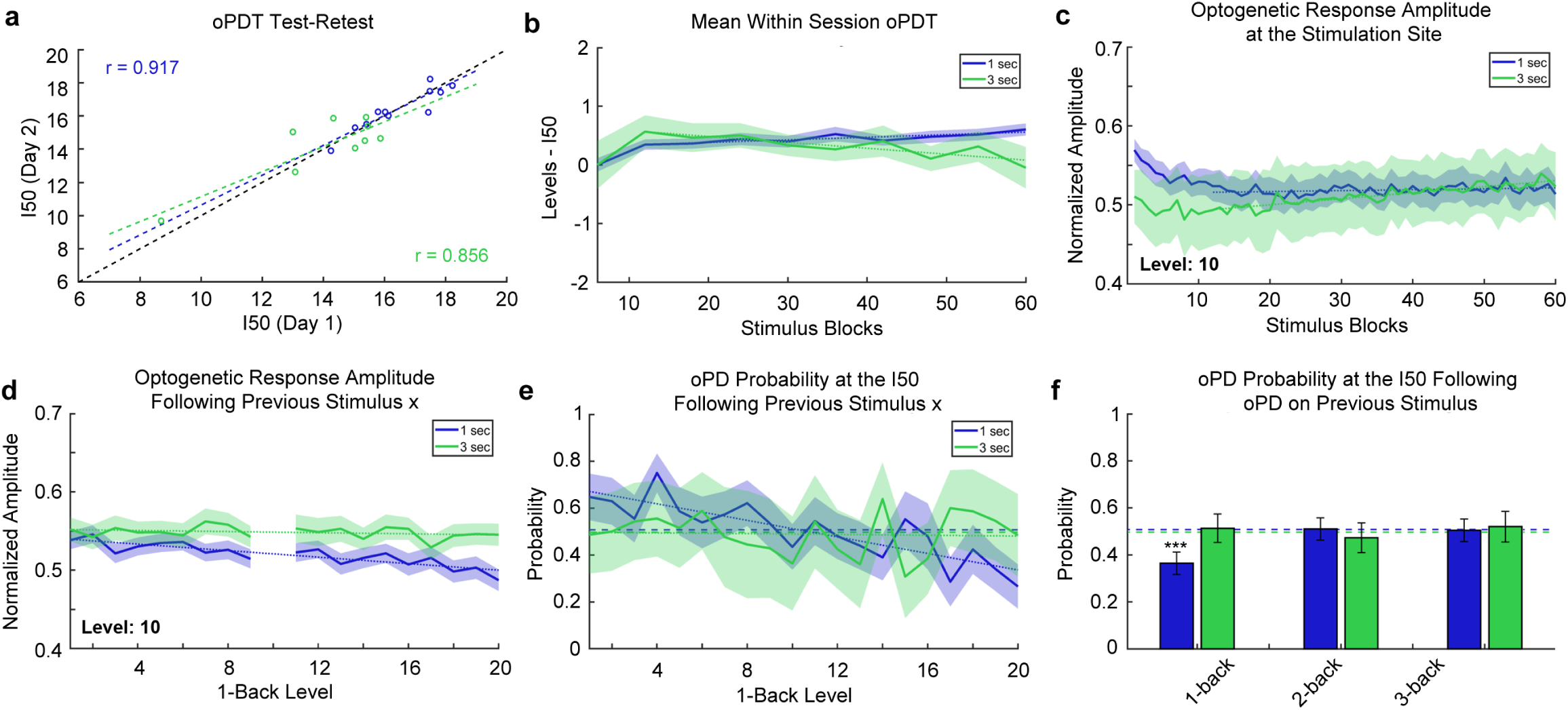
Stability and Reliability of the oPDT (**a**) Test-retest comparison of session I50s on consecutive days. n = 11 pairs for 1s int and n = 9 pairs for 3s int. Each session I50 is calculated from 60 replicates. Dashed black line indicates the diagonal. (**b**) I50 is relatively stable over the course of a session. Mean normalized intermediate I50s (6 replicate bins) for 1 and 3 sec ISIs. Each stimulus block consisted of 20 intensity levels presented in random order. Order was randomized independently for each block. Session duration is 20 min for the 1s ISI and 60 min for the 3s ISI. n = 67 and 41 sessions for the 1s and 3s ISI respectively. Shaded bars indicate 95% CI. Dotted lines indicate best fit (linear regression) excluding blocks 1-12. (**c**) Optogenetic response amplitude is reduced within the first 5 min of recording, likely due to ChR2 desensitization. Values were transformed via min-max normalization for the entire curve. Same dataset as in **b**. Dotted lines indicate best fit (linear regression) excluding blocks 1-12. (**d**) The optogenetic response amplitude at the stimulation site is reduced following high intensity stimuli (1-back stimuli are plotted on the x-axis). The y-axis indicates the mean normalized response amplitude at level 10. Same dataset as in **b**. (**e**) The probability of an oPD at the I50 depends on the intensity of the previous stimulus for 1s intervals. Dashed lines indicate the overall probability of an oPD at a level within ±0.25 of the I50. Dotted lines indicate best fit (linear regression). 1s: n = 29 sessions, 1740 trials, 3s: n = 10 sessions, 600 trials. (**f**) oPDs that occur < 2s prior have a suppressive effect on the probability of an oPD at the I50. Dashed lines indicate the overall probability of an oPD at a level within ±0.25 of the I50. Error bars indicate the 95% CI. Same subset as in **e**.

Mono-synaptic stimulation of CA1 afferents at 1 s intervals has been shown to induce long-term depression (LTD) *in vivo* (Thiels et al., 1994; Oliet et al., 1997; Massey and Bashir, 2007). In order to determine if the multi-synaptic I50 was drifting over time in our preparation, we broke each 60-block session into 10 x 6-block bins and calculated intermediate I50 values normalized by subtracting the overall session I50 (**Fig. 6b**). Following a transient increase for both, session I50 values gradually increased for the 1s interval and decreased for 3s interval (1s int: slope greater than zero, slope: 0.00442 [0.00206, 0.00678], p = 0.0031, 3s int: slope less than zero slope: −0.0101 [−0.0158, −0.00441], p = 0.0040, linear regression of the means excluding the first 12 replicates).

This gradual shift in the I50 through time could be a result of plasticity in the network but also ChR2 desensitization at the stimulation site. All ChR variants exhibit desensitization with varying degrees of reduction in peak photocurrents following repeated activation (Lin, 2010; Williams et al., 2013). We checked for ChR2 desensitization by measuring the peak amplitude (session min/max normalized) of the optogenetic response at the stimulation site over time (**Fig. 6c**). We found that the response amplitude is reduced (3%) within the first 4 minutes, likely an effect of desensitization, after which it increased slightly (1s int: slope greater than zero, slope: 1.16 x 10^−4^, 95% CI [7.62 x 10^−6^, 2.25 x 10^−4^], p = 0.037, 3s int: slope greater than zero, slope: 7.63 x 10^−4^, 95% CI [6.52 x 10^−4^, 8.75 x 10^−4^], p<0.0001, linear regression of the means excluding the first 12 replicates). Overall, the baseline drift in the response amplitude and the I50 over time was small relative to the effects of subthreshold PTZ, for instance (**Fig. 5b,f**). Nevertheless, drift must be accounted for in the interpretation of experimental results and all results should be compared to a vehicle control (**Fig. 5e,f**).

Next, we examined possible short term effects of stimulation on the P(oPD|I_x_) and assessed trial independence. The extent of ChR2 desensitization depends on the strength of previous activation and higher intensity stimuli are expected to produce greater desensitization than lower intensity stimuli. Since the stimuli are presented in random order, we were able to quantify this effect by measuring the extent of desensitization produced by each stimulus pair. We measured the peak amplitude of the response to a stimulus (level 10) following each of the other stimulus levels (**Fig. 6d**). There was a small reduction in amplitude of the response for strong preceding stimuli (1s int: slope less than zero, slope: −0.00208, 95% CI [−0.00271, −0.00145], p<0.0001, 3s int: slope not different from zero, slope: −3.58 x 10^−4^, 95% CI [−9.56 x 10^−4^, 2.40 x 10^−4^] p = 0.22, ns, linear regression of the means). In order to determine if this change in amplitude translated into a shift in the oPD probability, we calculated the oPDT probability at or near the I50 for each preceding stimulus (**Fig. 6e**). A subset of the data was selected for this analysis restricted to those sessions with I50 values that were near a presented stimulus level (within ± 0.25 levels of a whole number) (1s: *n* = 29 sessions, 1740 trials, 3s: *n* = 10 sessions, 600 trials). This allowed us to determine if deviations from chance were related to the stimulus order. The probability of oPD was negatively associated with the intensity of the previous stimulus for the 1s but not 3s interval (1s int: slope less than zero, slope: −0.0177, 95% CI [−0.0232, −0.0121], p<0.0001, 3s int: slope not different from zero, slope: −8.37 x 10^−4^ 95% CI [−0.008, 0.007], p = 0.82). This suggests that the reduction in amplitude (**Fig. 6d**) translates into a reduction in oPD probability.

Higher intensity stimuli are also more likely to produce an oPD, so we checked for effects of the oPD itself on the oPD probability at the I50. We calculated the conditional probability of an oPD at stimulus levels within ± 0.25 of the I50 (where the probability of an oPD should be close to chance), given an oPD on the nth previous stimulus (any level). Each n-back probability was calculated independently. We found a strong suppressive effect of a previous PD for the 1-back stimulus with a 1s ISI compared to the overall probability of an oPD at the I50 (p<0.0001, Fisher’s Exact Test, corrected for multiple comparisons using the Holm-Sidak method). PD probability did not differ significantly from chance at 2 and 3 back for the 1s interval (p = 0.91 and 0.32 respectively, Fisher’s Exact Test, corrected). PD probability remained at chance for 1, 2, and 3-back for the 3s interval (p = 0.91, 0.51, 0.45 respectively, Fisher’s Exact Test, corrected). Therefore, an interval of >2s is expected to be sufficient to produce independent trials and the 1s interval is expected to produce I50s that slightly overestimate the oPDT, likely due to a combination of ChR2 desensitization and after-hyperpolarization produced by the oPD. For this reason, it is important to compare measurements to baselines collected using the same protocol with the same ISI. It is important to note that all of the above measurements were taken from awake behaving animals and include random variation due to changes in ongoing activity. Taken together, these results indicate that the oPDT curve offers exceptional precision and reliability and as such provides a new clear window into the excitability state of the intact brain.

## Discussion

Neuronal activity occurs on a background of synaptic and intrinsic excitability governed by the biophysical properties of the component cells, synaptic plasticity, and the control of these parameters by internal homeostasis and brainstem modulatory systems (Sterling and Laughlin, 2015). The complex dynamics of neural circuitry makes it difficult to predict excitability from discrete measurements in reduced preparations, and obtaining direct measurements in behaving animals is technically challenging (Petersen, 2017). Changes in excitability relevant to seizure susceptibility are not fully captured by measurements of individual contributing factors and determining how these factors should be combined to define network excitability is a non-trivial problem. Using a novel light intensity-response procedure, we have developed and validated a new quantitative estimate of network excitability state based on PD probability. Network PDs are an unambiguous indicator of the transition from normal to abnormal functional states; by mapping the probability distribution of optogenetically induced PDs as a function of input magnitude, we derive a surrogate measure of network excitability, the I50, that depends not only on the excitability of the population of cells directly stimulated, but also on the receiving cells in the rest of the network. This approach has several important features: first, combining optogenetic modulators with a high precision LED light source allows for tighter control over induced currents in specific cell populations than previously possible with electrical stimulation; second, our multi-site recording technique allows us to simultaneously induce and monitor PDs as they propagate throughout the network, critical for unambiguous classification; third, our chronic preparation permits tracking of excitability dynamics over multiple timescales, from minutes to months; and finally, because the oPDT procedure does not produce seizures and the baseline I50 is relatively stable over time, pharmacological or molecular interventions can be tested within subjects, greatly increasing the ability to detect functional changes as a result of genetic and/or pharmacological manipulation.

The electrical ADT has been used extensively as a measure of seizure threshold (Löscher, 2017). In many ways, the oADT is the optogenetic equivalent of the electrical ADT, both produce similar acute after-discharges and repeated presentations lead to behavioral seizure (kindling). Although seizures are an unmistakable sign that a threshold has been crossed, they also produce long lasting effects on the brain, including widespread changes in gene expression (Altar et al., 2004). In contrast, the oPDT does not produce seizures, nor kindling, and therefore can be used to measure thresholds over time allowing within-subjects comparisons free from the confounds of seizure exposure. This property may allow the oPDT to be used to track changes in excitability in models of epileptogenesis as a way to gain insight into mechanisms and measure the effectiveness of various pharmacological and molecular interventions.

Previously optogenetic seizure thresholds have been determined by varying the number of stimulus bouts (Khoshkhoo 2017). The oADT represents the first example of systematic optogenetic intensity-response curves, using a high-precision LED, to precisely determine seizure thresholds (keeping stimulation frequency, pulse width, and train duration constant). When the oADT and the oPDT are combined, the oADT, in the context of a known oPDT provides additional information about seizure susceptibility. Although the thresholds for both metrics are the same in terms of light intensity, the oADT requires repeated oPD inductions in order to produce an AD. This suggests that distinct mechanisms are involved in the oAD generation, possibly the accumulation of extracellular K+ and the switch to depolarizing GABA (Alfonsa et al., 2015; Buchin et al., 2016; Cossart et al., 2005; Miles et al., 2012). Comparing the results of each approach might allow for identification of drugs and manipulations that prevent AD but do not change baseline excitability, and therefore can be expected to have fewer side effects.

Electrically-evoked potential amplitude has previously been used as a measure of excitability in animal models and patients with epilepsy (Maru and Goddard, 1987; Freestone et al., 2011; Enatsu et al., 2012; Wendling et al., 2016; Keller et al., 2018). However, without knowing the details of underlying synaptic currents and their influence on the LFP, it is difficult to estimate excitability from amplitude alone given that somatic inhibition and dendritic excitation produce similar sink/source patterns (Buzsáki et al., 2012; Hales and Pockett, 2014; Herreras, 2016). Because network wide PDs are readily differentiated from subthreshold synaptic potentials, our preparation provides an unambiguous measurement of excitability that does not require knowledge of the sink/source patterns needed to interpret the LFP itself.

Electrical stimulation activates cells and fibers of passage which act upon a sparse and widely distributed population of neurons whose identity cannot be predicted ahead of time, nor known with certainty afterwards (Histed et al., 2009). In contrast, optogenetic expression patterns can readily be determined histologically and light spread can be accurately modeled based on fiber location and power output. Light spreads out in a cone from the tip of the fiber and drops off quickly with distance in brain tissue due to scattering (Aravanis et al., 2007; Yizhar et al., 2011). As the light intensity increases, cells directly under the fiber will be increasingly depolarized to threshold followed by neighboring cells. Thus, the optogenetic intensity-response curve varies both the magnitude of depolarization in individual cells as well as the overall size of the activated population. Cells in the center that have already reached threshold cannot be depolarized further, in part because ChR2 currents are voltage dependent (Williams et al., 2013). From the perspective of downstream areas, the intensity-response curve varies the number of synchronously active projections emanating from the stimulation site. This is important because, while synchronous firing is an important feature of the function of these circuits, exceeding normal ranges would be expected to produce non-specific propagation of the synchronous discharge when some critical number of downstream cells is induced to fire synchronously.

The sharp transition between sub and suprathreshold activity may reflect a breakdown between the balance of feedforward inhibition and excitation (Buzsáki, 1984; Wahlstrom-Helgren and Klyachko, 2016). If the magnitude of the population EPSP exceeds that of feedforward inhibition, it could produce a highly synchronous discharge that would propagate to other areas (Johnston and Brown, 1981). This would explain the sensitivity of the oPDT curve to GABA antagonists such as PTZ. In addition to the balance of inhibition and excitation, a number of other functional properties might be expected to modulate the oPDT, the oADT, or both. Synaptic strength determines the threshold for downstream propagation of the synchronous discharge (Nicoll, 2017), intrinsic excitability modulates the responsiveness of downstream cells, which is governed by the relative expression and cellular localization of voltage-gated Na^+^, K^+^, and Ca^2+^ channels (Graef and Godwin, 2010; Lisman et al., 2018; Meadows et al., 2016), and the regulation of the extracellular space by glia (Devinsky et al., 2013).

The Thy1-ChR2 mice (line 18) used in these experiments express high levels of wild-type ChR2-EYFP in deep (calbindin-negative) pyramidal neurons in CA1 (Arenkiel et al., 2007; Dobbins et al., 2018). Importantly, transgenic animals provide more consistent and specific ChR2 expression patterns than could be achieved with virus injection. Compared to neighboring superficial cells, deep pyramidal neurons in CA1 are more prone to burst firing, have unique input and output pathways, and lack calcium buffering proteins, potentially making them more susceptible to seizure activity (Mizuseki et al., 2011; Sloviter, 1989; Valero et al., 2015). By selectively targeting this population of excitatory neurons the inhibitory population is free to respond, preserving the strong reciprocal relationship they have with the excitatory cells being stimulated, and allowing them to participate in the generation of the synchronous discharge (Cobb et al., 1995; Ellender et al., 2014; Sessolo et al., 2015; Yekhlef et al., 2015; Khoshkhoo et al., 2017; Wang et al., 2017; Chang et al., 2018a; Magloire et al., 2018).

The fact that the evoked discharges initiated in CA1 occurred simultaneously in multiple hippocampal structures supports the hypothesis that synchronous activation of entorhinal cortex was responsible (Avoli et al., 2002; Yeckel and Berger, 1990). The oPD was most prominent in areas CA3 and DG; reciprocally connected areas prone to hypersynchronous discharge (Traub and Jefferys, 1994). CA3 and DG lead during spontaneous discharges suggesting that synchronous inputs to these areas are responsible for generating the IIS, and optogenetically generated synchronous input from CA1 via subiculum and entorhinal cortex serves as a trigger for the oPD. These results are also consistent with the dentate gate hypothesis, which posits that dentate granule cells, because of their relatively high functional threshold, act as a gate that prevents the propagation of seizure activity into the rest of hippocampus (Heinemann et al., 1992; Lothman et al., 1992; Goldberg and Coulter, 2013). Although hippocampus has multiple interacting nested loops, if the dentate gyrus is the last to reach “critical mass”, a synchronous discharge there could be responsible for the all-or-none expression of the oPD. It this is true, the oPDT, in its current configuration, is a sensitive measure of the strength of the “dentate gate” and could be useful for evaluating therapeutic interventions in this area (Krook-Magnuson et al., 2013, 2015). Non-synaptic propagation, often neglected in earlier studies of seizure, is also likely to play a role in generating the synchronous discharge in the closely situated structures of the hippocampus (Zhang et al., 2014).

Despite the widespread synchronous nature of the discharge, it is transient, and the system is surprisingly resilient to its effects on short time scales. By analyzing the oPD probabilities from one stimulus to the next, we found that the response is functionally independent of the previous stimulus in just a few seconds. Furthermore, closely spaced oPDs (1s ISI) are suppressive to the system, significantly reducing the probability of another oPD in response to the subsequent stimulus. The suppressive effect of the oPD is likely due to a combination of ChR2 desensitization and the slow after-hyperpolarization (I_sAHP_) (Alger and Nicoll, 1980; Lancaster and Adams, 1986). The fact that unitary PDs are suppressive and multiple repeated oPDs are required to initiate seizure suggests that either there is a reduction in the suppressive effect with each oPD, or that multiple PDs produce sufficient excitation to overcome the suppressive effects. In the context of spontaneous seizure, this makes it clear that seizures are a result of a slow transition to abnormal brain states, such as those presumably induced by persistence repetitive synchronous discharges in our model, which are permissive to self-sustaining seizure activity and that they are not simply the result of transient synchronous activity alone (Chang et al., 2018b).

The oPDT procedure allows for multiple threshold measurements over time; an important advantage over previous methods. For comparison, the recovery time for the 6hz – psychomotor seizure and minimal electroshock threshold is at least 4.4 and 7.5 minutes respectively (Brown et al., 1953). The enhanced temporal resolution of the oPDT can be used to estimate the functional pharmacokinetics of anti-epileptic drugs or to study the natural fluctuations in excitability that depend on brain state and behavior (Baud et al., 2018, 2019). The temporal resolution of the oPDT is limited by the number of stimuli used to generate the curve and the ISI. As we have demonstrated, stimuli are functionally independent when presented with ISI > 2s. Further reductions could be achieved by using a reduced subset of stimuli, optimized for baseline curves in each animal. For small changes, sampling at the I50 alone should be sufficient. When implemented in a closed-loop system, optimization techniques could be used to further reduce the number of stimuli necessary (MacKay, 1992).

On longer time scales (days to weeks), repeated stimulation of the brain is expected to produce a number of changes, some of which involve changes in plasticity and homeostatic regulation of the stimulated cells (Thiels et al., 1994; Oliet et al., 1997; Massey and Bashir, 2007). Recently, it was demonstrated that long-term (24hrs) optogenetic stimulation of CA1 produces changes in spine density and synaptic plasticity (Moulin et al., 2019). Although we have demonstrated that the oPDT is stable over time, appropriate control experiments should be used and care should be taken when interpreting results to account for potential changes in the brain as a result of stimulation.

The current study is focused on using optogenetic intensity-response curves to quantify excitability in the context of seizure activity, but the same procedure could be applied to any study that uses optogenetic stimulation as the independent variable with a stochastic dependent variable. Similar procedures could be performed with any appropriate optogenetically induced cellular, molecular, physiological, or behavioral output metric (Allen et al., 2015; Bugaj et al., 2016; Tischer and Weiner, 2014). Abnormal excitability is also associated with a number of other neurological disorders including depression, autism, Alzheimer’s, anxiety, drug addiction (and withdrawal), and schizophrenia (Crabtree et al., 2017; Friedman et al., 2014; Kourrich et al., 2015; Rau et al., 2015; Santos et al., 2010; Takarae and Sweeney, 2017). Correlating changes in excitability with behavior or neurochemical state in the context of these disorders may provide novel insights into the mechanisms of the underlying disease.

In summary, we show that synchronous optogenetic activation is sufficient to induce epileptiform population discharges, and the systematic modulation of light intensity allows for precise estimation of the threshold between normal and abnormal activity. This approach provides a novel platform for testing the effects of therapeutic interventions on network excitability.

## Acknowledgements

Special thanks to: Dorothy Dobbins, Allison Goldstein, Rasesh Joshi, Glen Marrs, and Hong Qu Shan for help with this project and useful comments on the manuscript. Funding was provided by the NIH grants R01AA016852, T32AA007565, F31AA021322, and the Tab Williams Family Fund. Portions of this work were previously reported in abstract form (Klorig et al., 2014, 2017).

## Contributions

DK and GA designed experiments and performed surgeries. DK, GA, and TS collected data. DK developed the analysis. DK, DG, and GA wrote the paper.

