## Supplemental Figures for "Optogenetically-Induced Population Discharge Threshold as a Sensitive Measure of Network Excitability"

### oPDT Supplementary Experimental Procedures and Figures

### S1: Spontaneous and Evoked Population Discharges

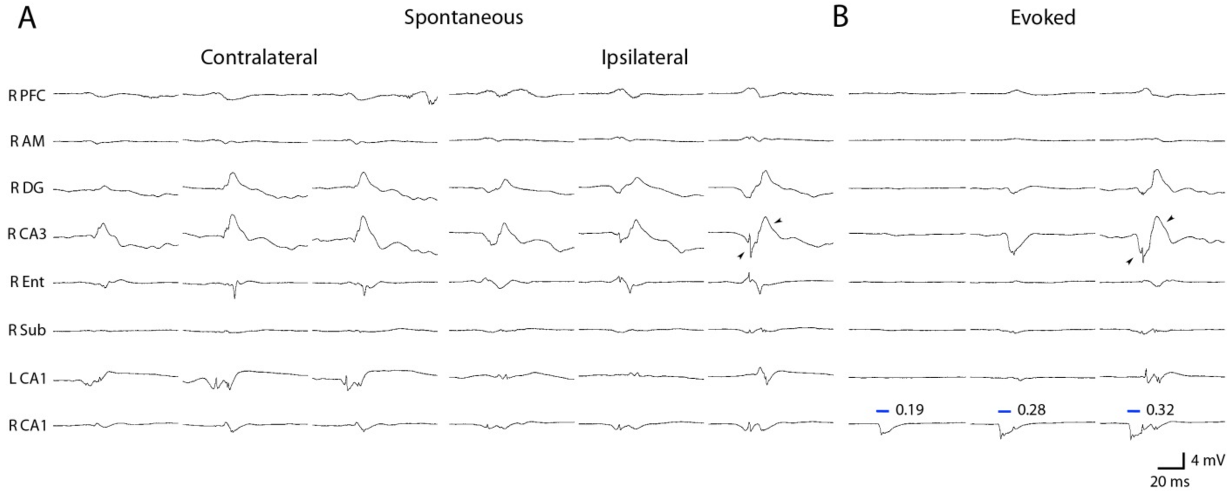


**Figure S1: Optogenetic Stimulation Evokes Network Wide Population Discharges that Resemble Spontaneous Interictal Spikes**

A. Comparison of spontaneous interictal discharges to optogenetically evoked PDs. Spontaneous contralateral spikes are characterized by leading activity in contralateral (left) CA1. Note the positive going (presumably inhibitory) wave in DG and CA3. Ipsilateral spikes are characterized by near synchronous activation in DG, CA3, and R CA1, and delayed activation in L CA1. B. Evoked population spikes are led by short latency activation in CA1 (stim site) followed by near synchronous activation of DG, CA3, and CA1 with a latency of ~10 ms. Note similar timing and wave shape between post-stimulus evoked population spikes and spontaneous ipsilateral spikes (arrow heads). Blue bars indicate onset and duration of light stimulus in CA1.

S2: Electrode Placement

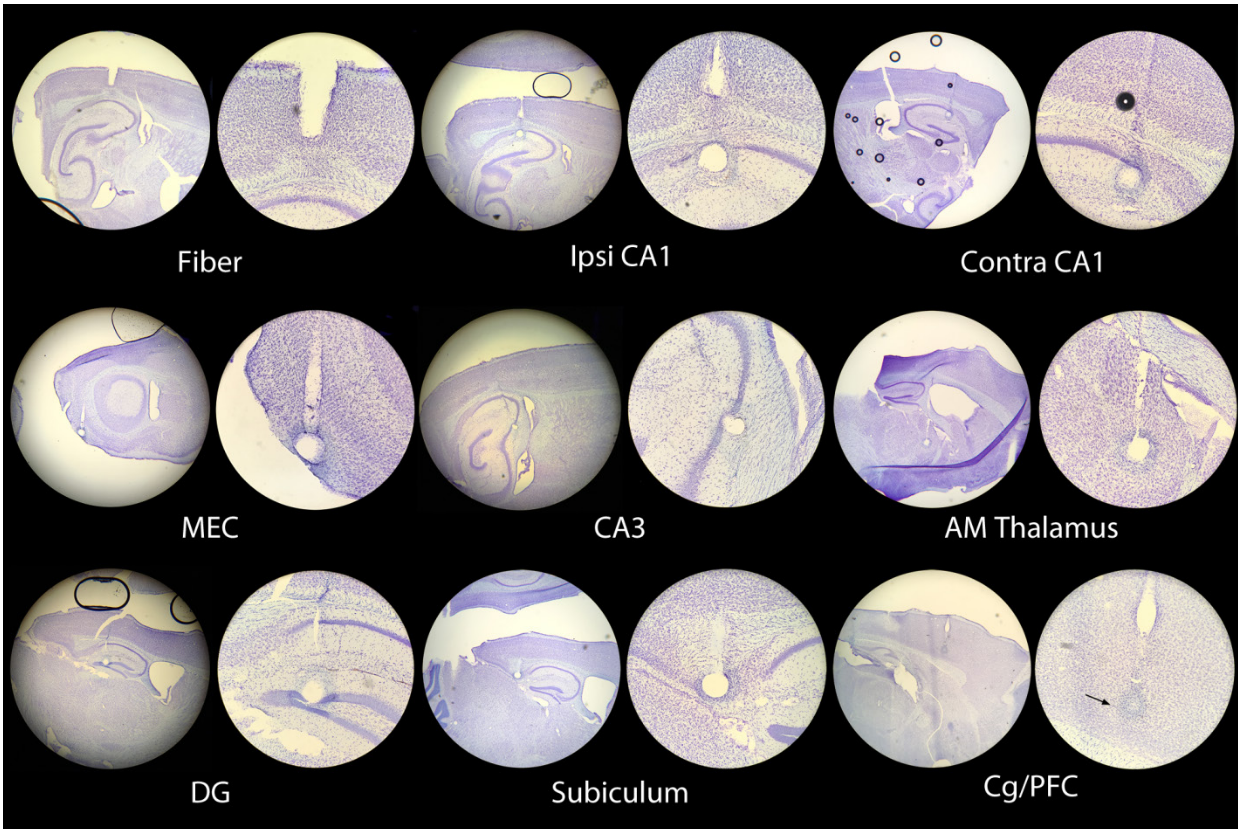


**Supplementary Figure S2: Satellite array electrode placement.** Electrolytic lesions were performed just prior to perfusion on each animal to help localize the electrode tip. Nissl stain.

**S3: Behavioral Seizure Staging**

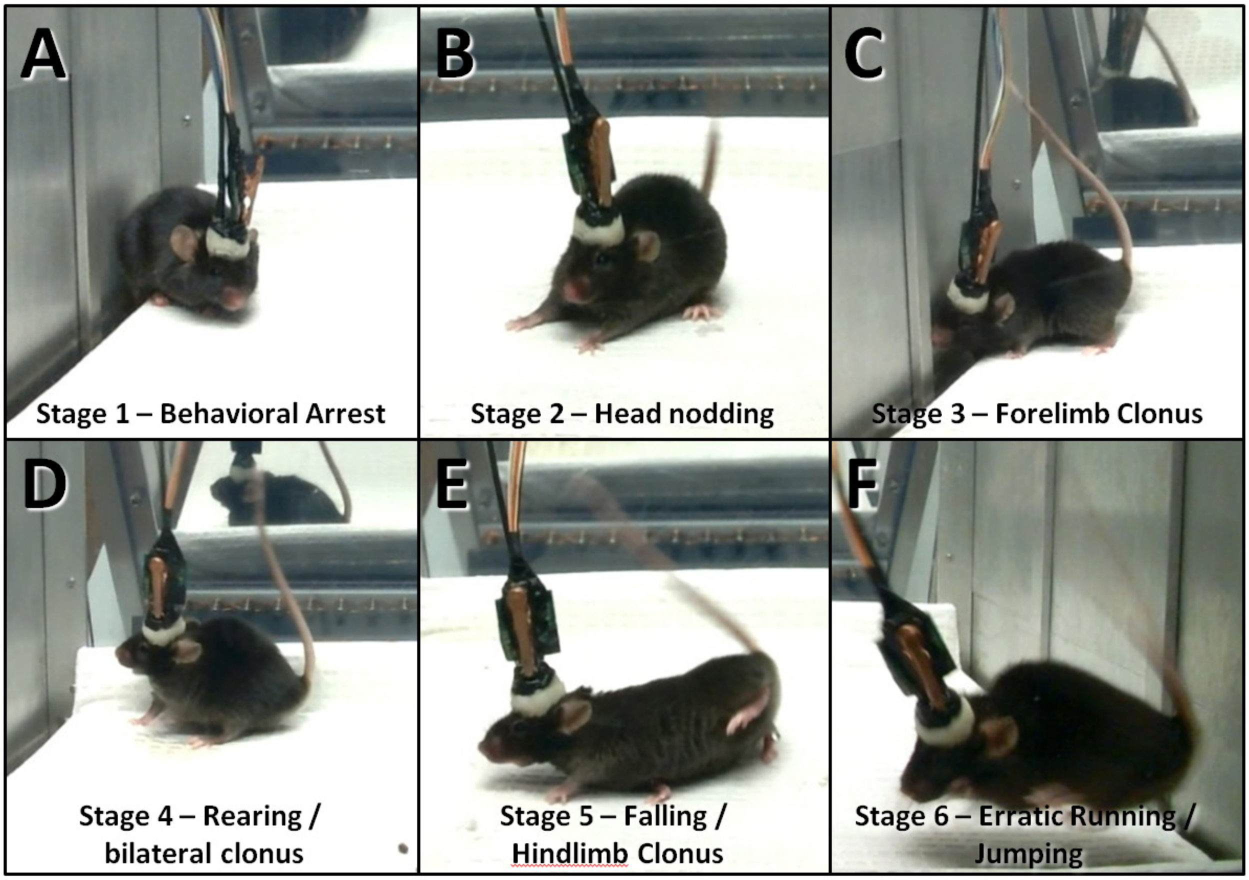


**Supplementary Figure S3. Behavioral Manifestations in Optogenetically Kindled Mice.** A modified Racine scale was used to quantify seizure severity.

**S4: Raw Traces Depicting oPD**

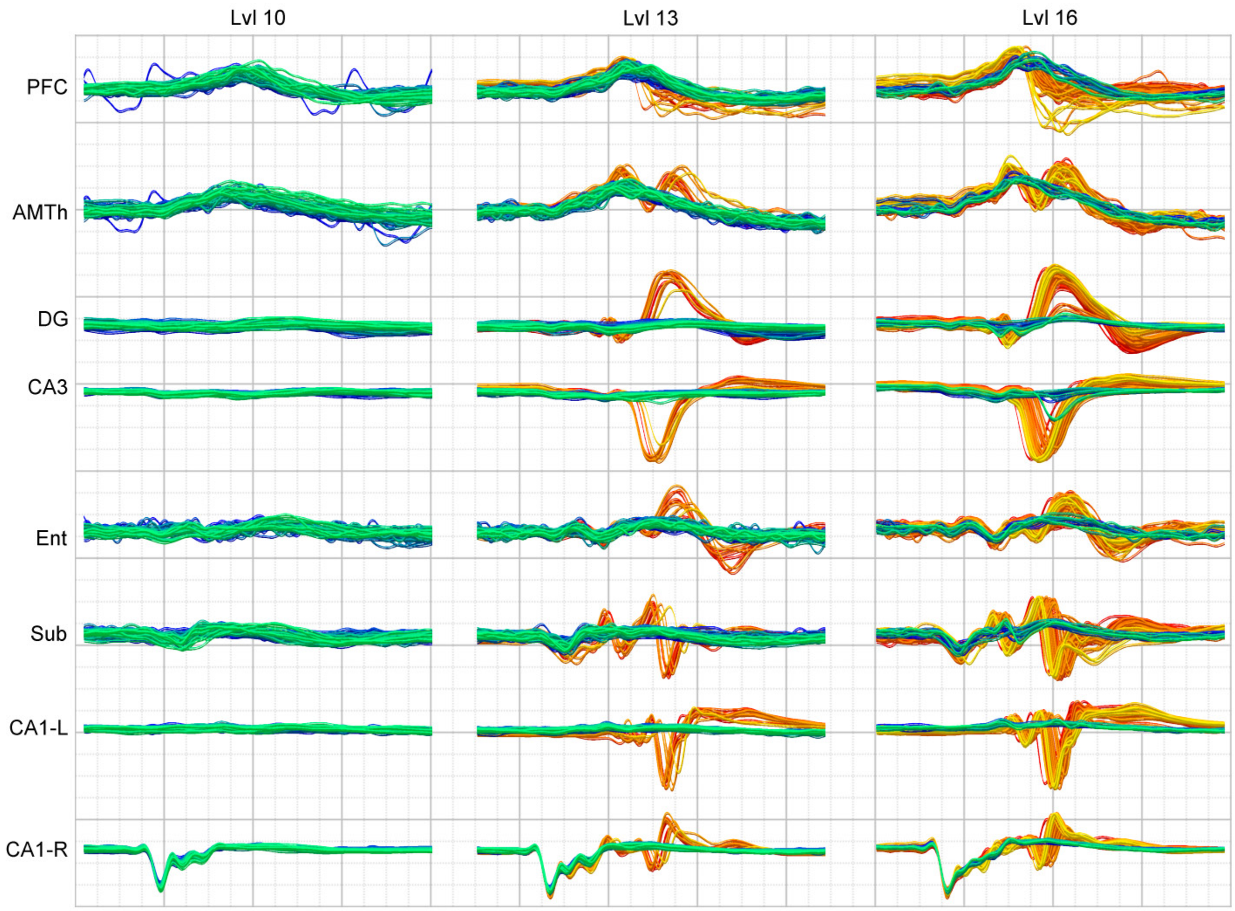


**Supplementary Figure S4. Raw Traces Depicting oPD.** Traces are scaled by the maximum amplitude response and color coded by state. Each trace is 200 ms long, -50 ms pre, +150 post stim.

### S5: Light Dose-Response Bistability

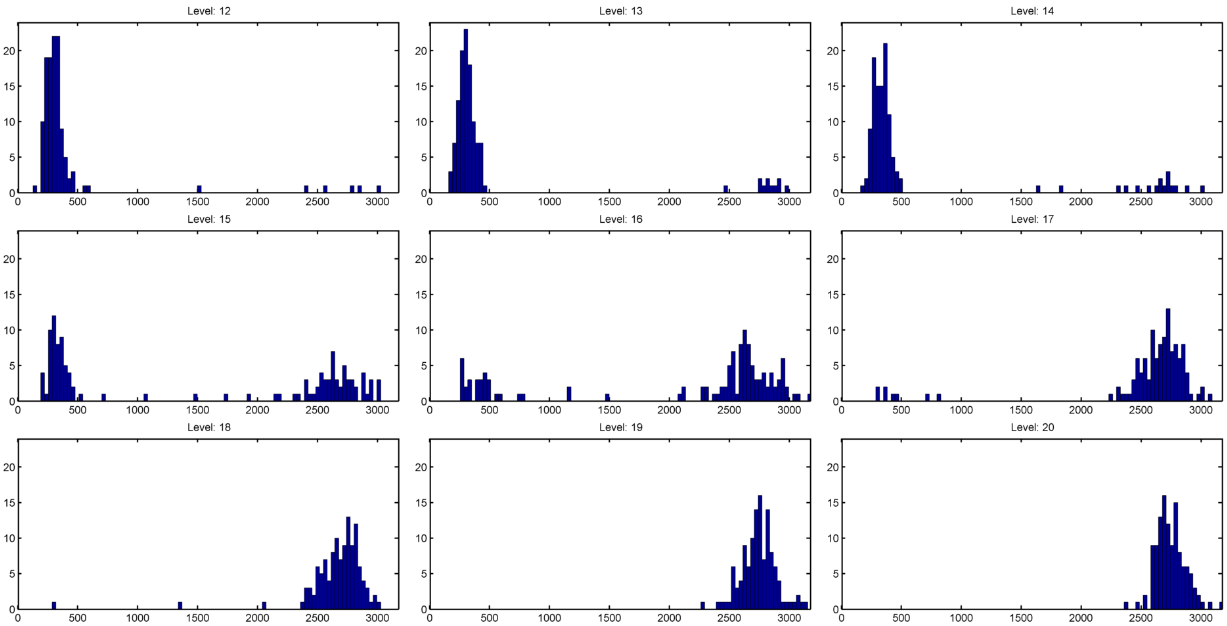


**Supplementary Figure S5. Light Dose Response Histogram.** Example histograms of maximum response amplitude (max-min) at each intensity level (same data as depicted in figure 3e-h). Y is counts, x is evoked LFP amplitude (n = 120 reps).
